# Durable protection against SIV challenge by adeno-associated virus delivery of Env-specific antibodies

**DOI:** 10.64898/2026.04.20.719478

**Authors:** Natasha M. Clark, Nida K. Keles, Grace N. Hagedorn, Vadim A. Klenchin, Hanne Andersen, Jim Treece, Christine M. Fennessey, Jun Xie, Thomas N. Denny, C. Todd DeMarco, Jose S. Hernandez, Saverio Capuano, Matthew R. Gardner, Brandon F. Keele, Guangping Gao, Mark G. Lewis, Mario Roederer, David T. Evans

## Abstract

Conventional vaccines have so far failed to elicit the types of antibodies needed for protection against HIV. As an alternative, we evaluated adeno-associated virus (AAV) delivery of rhesus macaque antibodies to the SIV envelope glycoprotein for protection against SIV challenge. AAV vectors encoding a broadly neutralizing antibody (bnAb) and an antibody that only mediates antibody-dependent cellular cytotoxicity (ADCC) were administered individually or together to separate groups of rhesus macaques. Antibody expression was sustained for more than a year with minimal anti-drug antibody responses. All animals that received a control antibody or the ADCC-only antibody became infected after five low-dose, intrarectal challenges with SIV_mac_239. In contrast, 14 of 16 animals that received the bnAb resisted two rounds of twelve SIV_mac_239 challenges more than a year apart. Thus, AAV delivery of a single bnAb can afford durable protection against a pathogenic SIV strain that is notoriously difficult to protect against by vaccination.

## INTRODUCTION

The development of a safe and effective vaccine to curb the global spread of HIV remains one of the greatest scientific challenges of our time. Dozens of potent, broadly neutralizing antibodies (bnAbs) targeting conserved, conformational features of the HIV envelope glycoprotein (Env) have been isolated in recent years. Passive transfer of many of these bnAbs have afforded protection against simian-human immunodeficiency virus (SHIV) challenge in nonhuman primate models (*1–3*) and against HIV acquisition in one clinical trial (*4, 5*). However, efforts to elicit these types of antibodies by vaccination have so far been unsuccessful. Practical limitations also preclude repeated infusion of bnAbs to maintain protective antibody concentrations in individuals at risk of HIV infection.

Transduction of muscle cells with adeno-associated virus (AAV) vectors encoding bnAbs can afford long-term expression of antibodies that protect against HIV in humanized mice (*6, 7*), and against SHIV if administered to neonatal macaques (*8*). Although antibodies to the vectored antibodies, termed anti-drug antibodies (ADAs), severely impair bnAb expression in most animals (*9, 10*), we recently demonstrated sustained expression of natural, species-matched antibodies with minimal ADAs in immunocompetent adult rhesus macaques followed by containment of SIV rebound after discontinuing antiretroviral drug therapy (*11*). In the present study, we show that AAV delivery of antibodies with potent neutralization and ADCC can afford complete protection against mucosal challenge with SIV_mac_239 for at least a year after AAV administration. SIV_mac_239 is a tier 3 neutralization-resistant virus that is particularly well-adapted for replication in rhesus macaques. These results therefore provide a rigorous illustration of the potential for AAV delivery of bnAbs to provide lasting protection against HIV acquisition in people.

## RESULTS

### AAV delivery of SIV Env-specific antibodies

AAV vectors encoding two rhesus macaque antibodies to the SIV envelope glycoprotein, ITS61.01 and ITS103.01 (*12*), and a control antibody to the respiratory syncytial virus fusion protein, 17-HD9 (*13*), were engineered as previously described (*11*). ITS61.01 binds to the gp120 high mannose patch (*12*) and ITS103.01 binds to a conserved epitope in the CD4 binding site (*14*). Both antibodies direct efficient ADCC against SIV_mac_239-infected cells (*15*). However, only ITS103.01 potently neutralizes SIV_mac_239 infectivity (*15*).

Thirty-two adult rhesus macaques of Indian ancestry were selected for this study after pre-screening to exclude animals with neutralizing antibody titers to AAV serotype 9 (AAV9) (*16*) and MHC class I alleles associated with spontaneous control of SIV replication (*Mamu-B*008* and *-B*017* (*17, 18*)). AAV9 packaged vectors encoding ITS61.01 and ITS103.01 were administered alone or in combination to three separate groups of eight animals by intramuscular injection (3.3×10^12^ gc/kg each). Eight additional control animals received an AAV9 vector encoding 17-HD9.

Four weeks after AAV delivery, both Env-specific antibodies were detectable in serum from all treated animals but became more variable by week 8. By week 16, serum concentrations ranged from 0 to 146.2 (mean 45.2) µg/ml for ITS61.01 and from 0 to 137.9 (mean 46.5) µg/ml for ITS103.01 (Fig. 1A, B and Table S1). ITS61.01 and ITS103.01 were also detectable in rectal transudate at weeks 8 and 12 post-AAV administration at concentrations that correlated with serum concentrations at these timepoints (Fig. S1A-D).

**Fig. 1.**
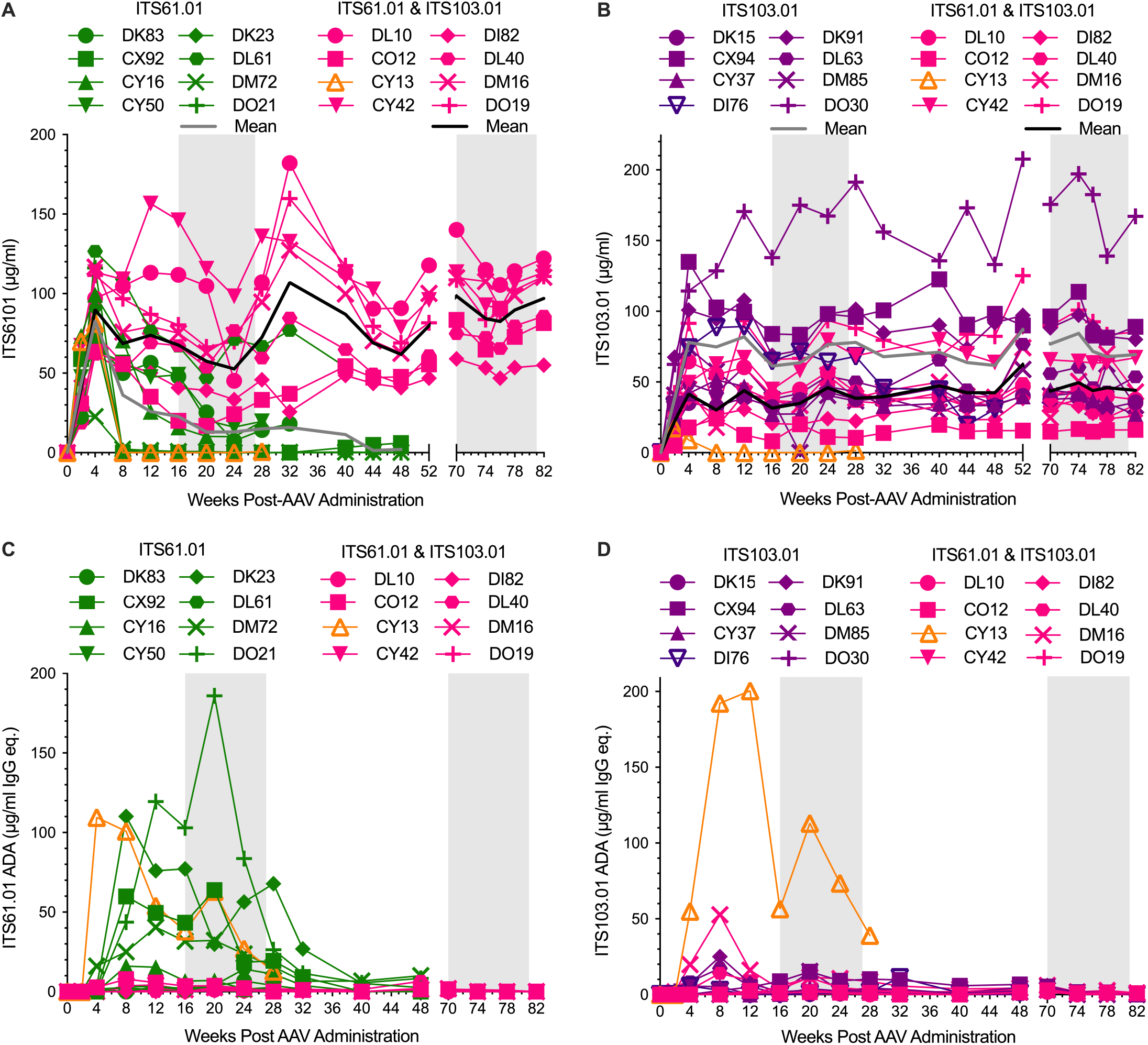
Serum antibody concentrations and anti-drug antibody responses. ITS61.01 (**A**) and ITS103.01 (**B**) concentrations in serum were measured by ELISA on plates coated with an antibody to a rhodopsin tag appended to the C-terminus of ITS61.01 or with an anti-idiotype antibody to ITS103.01. The mean antibody concentrations in each of the treatment groups (N=8) is indicated by the grey and black lines. Anti-drug antibody responses to ITS61.01 (**C**) and ITS103.01 (**D**) were measured by probing ITS61.01- or ITS103.01-coated plates with biotinylated IgG purified from serum followed by detection with streptavidin-HRP and development with TMB substrate. Plotted values represent the average of absorbance reads across all dilutions falling within the range of a valid calibration curve, in most cases 3-4 technical replicates. Symbols and lines are color-coded to indicate the animals that received vectors encoding ITS61.01 (green), ITS103.01 (purple and indigo) or ITS61.01 & ITS103.01 (light red and orange). The shaded regions indicate twelve-week periods of repeated, low dose intrarectal challenge with SIV_mac_239.

ADA responses were generally low and reflected an inverse relationship with ITS61.01 and ITS103.01. The only animal with consistently high ADA titers to both ITS61.01 and ITS103.01 (CY13) had negligible serum concentrations of both antibodies by week 8 post-AAV administration (Fig. 1C and D). Four of the animals that received the vector encoding ITS61.01 alone also had high ADA titers (Fig. 1C), which accounts for the inability of these animals to maintain detectable levels of this antibody.

### Protection against mucosal challenge with SIV_mac_239

Sixteen weeks after AAV9 administration, the macaques were exposed to twelve weekly low dose (150 TCID_50_), intrarectal challenges with SIV_mac_239. All animals that received vectors encoding the control antibody 17-HD9 or the ADCC-only antibody ITS61.01 became infected after five challenges (Fig. 2A). In contrast, fourteen of the sixteen animals that received the vector encoding the bnAb ITS103.01, either alone or in combination with the vector for ITS61.01, resisted all twelve challenges (Fig. 2A). Differences in the rate of SIV infection between the groups that received ITS103.01 and the control antibody were highly significant (Fig. 2B).

**Fig. 2.**
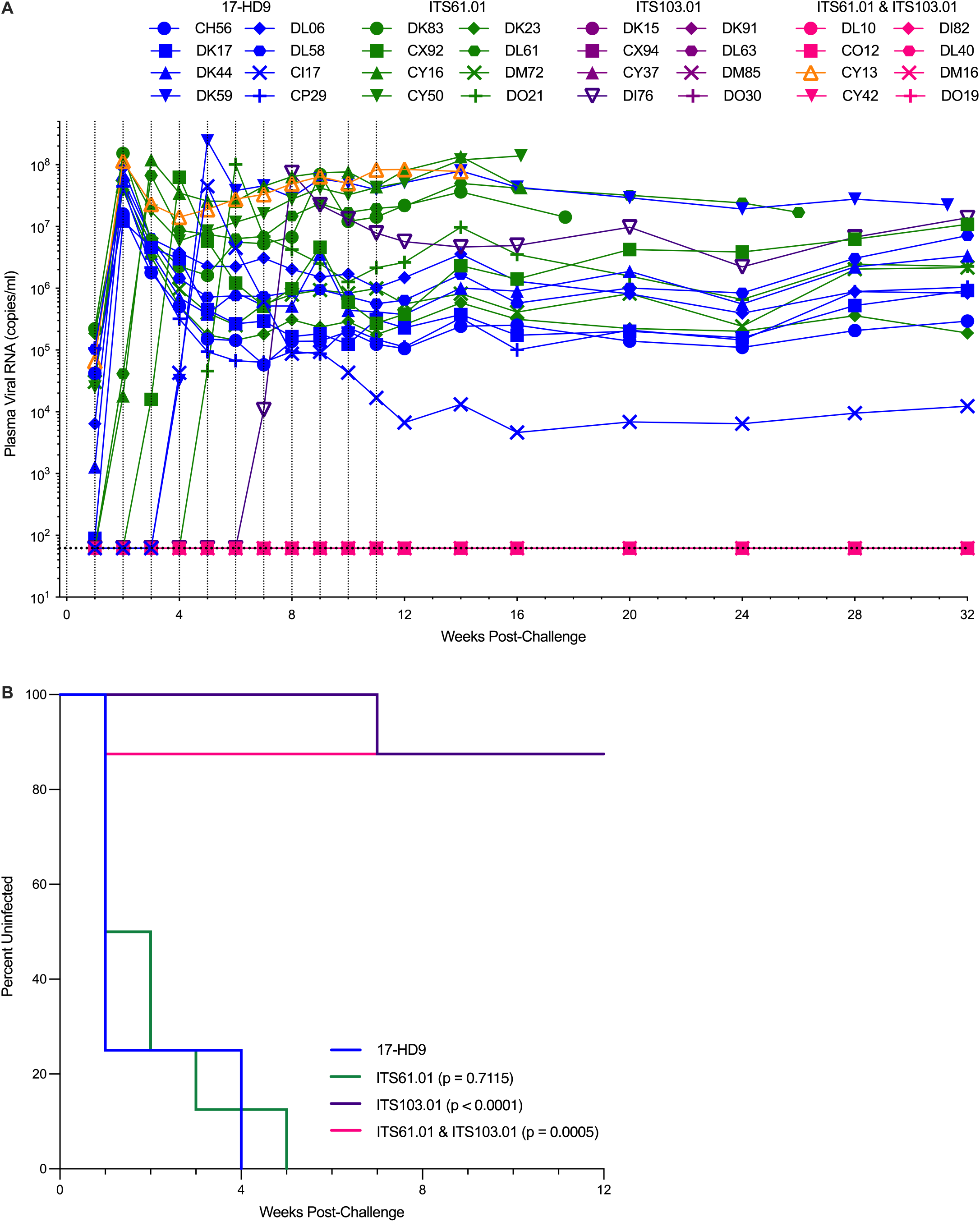
Outcome of SIV challenge beginning at week 16 post-AAV. (**A**) SIV RNA loads in plasma were measured using a qRT-PCR assay with a quantification threshold of 62 copies/ml (dotted line). Plotted values represent the mean of two replicate reactions per sample. The vertical dotted lines indicate the twelve intrarectal challenges with SIV_mac_239 (150 TCID_50_). (**B**) Differences in the percentage of uninfected animals remaining over the course of SIV challenge for groups that received Env-specific antibodies (N=8 each) were compared to the group that received the control antibody (N=8) using the Mantel-Cox test (ITS61.01 p=0.7115, ITS103.01 p<0.0001 and ITS61.01 & ITS103.01 p=0.0005). Symbols and lines are color-coded to indicate the animals that received vectors encoding 17-HD9 (blue), ITS61.01 (green), ITS103.01 (purple and indigo) or ITS61.01 & ITS103.01 (light red and orange).

Of the two animals that became infected with SIV despite AAV delivery of ITS103.01, one received vectors for both ITS103.01 and ITS61.01 (CY13) and the other received the vector for ITS103.01 alone (DI76). CY13 became infected after the first SIV challenge in the absence of detectable concentrations of either antibody owing to high ADA levels, whereas DI76 became infected after the seventh challenge despite an above average serum concentration of ITS103.01 (∼65 µg/ml). Sequence analysis of the virus population in plasma did not reveal any consistent changes in *env* among the animals expressing only ITS61.01 or in animal CY13 (Fig. S2A).

However, Env sequences from DI76 had an N479D substitution predicted to eliminate an N-linked glycan flanking the CD4-binding site (Fig. S2B), which was previously shown to confer complete resistance to neutralization and ADCC by ITS103.01 (*11*). Thus, failure to protect against SIV challenge can be explained by high ADA responses in CY13 and antibody escape in DI76.

More than a year after AAV delivery, serum concentrations of ITS61.01 and ITS103.01 in the protected animals remained high and unabated (Fig. 1A, B and Table S2). Both antibodies were also detectable in rectal transudate 69 weeks after AAV administration (Fig. S1E, F), confirming their continued presence at the site of challenge. Beginning at week 70, these animals were therefore rechallenged alongside six naïve control animals with another twelve weekly intrarectal doses of SIV_mac_239 (150 TCID_50_). While the control animals all became infected after six challenges, the animals that received ITS103.01 resisted all twelve challenges (Fig. 3A).

**Fig. 3.**
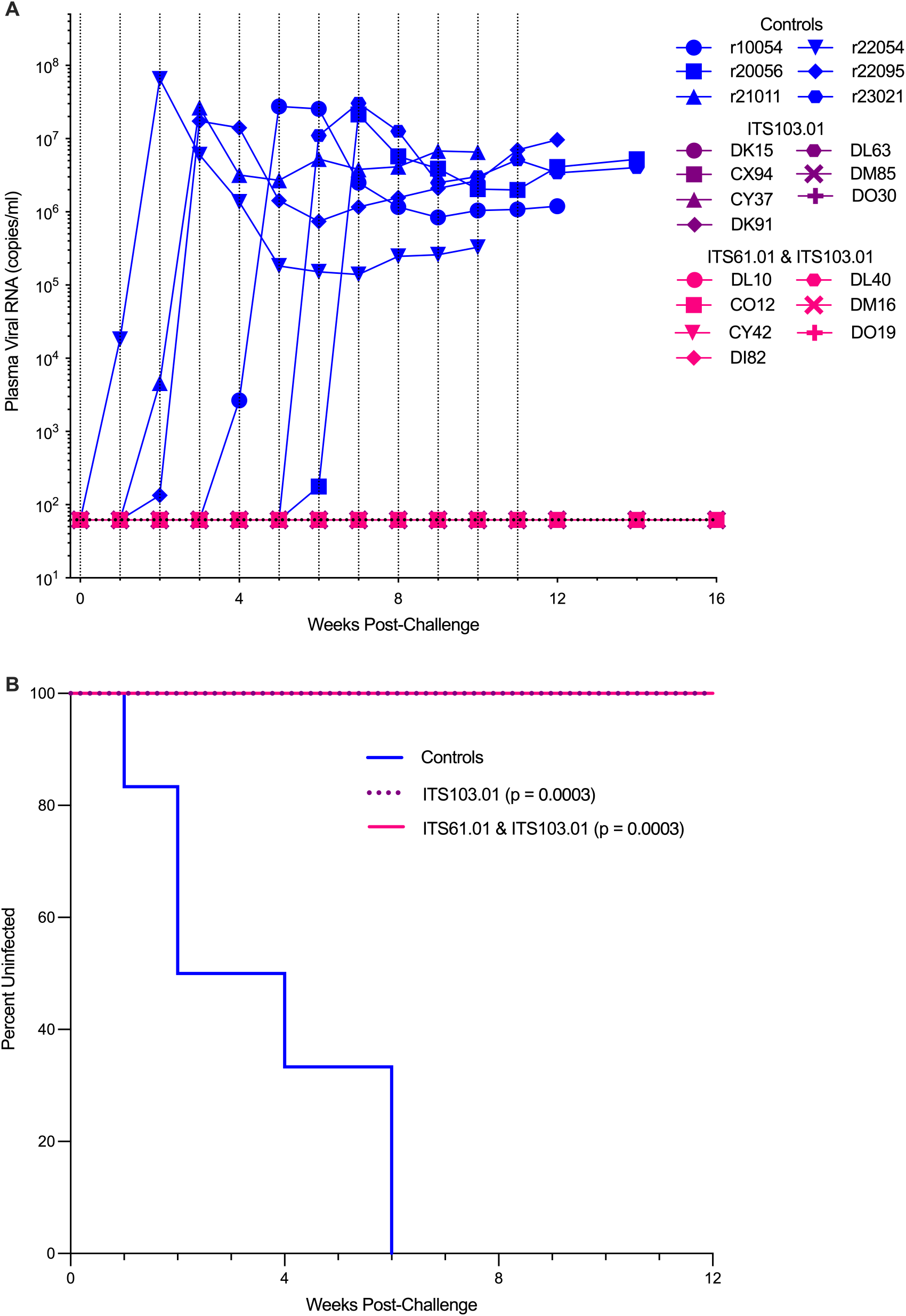
Outcome of SIV challenge beginning at week 70 post-AAV. (**A**) SIV RNA loads in plasma were measured using a qRT-PCR assay with a quantification threshold of 62 copies/ml (dotted line). Plotted values represent the mean of two replicate reactions per sample. The vertical dotted lines indicate the twelve intrarectal challenges with SIV_mac_239 (150 TCID_50_). (**B**) Differences in the percentage of uninfected animals remaining over the course of SIV challenge for the groups that received Env-specific antibodies (N=7 each) were compared to the naïve control group (N=6) using the Mantel-Cox test (ITS103.01 p=0.0003 and ITS61.01 & ITS103.01 p=0.0003). Symbols and lines are color-coded to indicate the naïve control animals (blue) and the animals that received vectors encoding ITS103.01 (purple) and ITS61.01 & ITS103.01 (light red).

Differences in the rate of infection between the ITS103.01 groups and the control group were again highly significant (Fig. 3B). Thus, AAV delivery of ITS103.01 provided complete protection against SIV_mac_239 rechallenge for well over a year after AAV administration.

### SIV_mac_239 neutralization and ADCC

Sera collected at six-week intervals beginning at the time of the first SIV challenge (weeks 16, 22 and 28 post-AAV) and at the time of rechallenge (weeks 70, 76 and 82 post-AAV) were tested for neutralization of SIV_mac_239. Samples from the animals that received ITS103.01 alone or together with ITS61.01 potently neutralized SIV_mac_239 at all time points, whereas only low titer endogenous neutralizing antibodies were observed in three ITS61.01-only animals after they became infected with SIV (Fig. 4A, B). During the initial challenge phase, IC50 titers ranged from 2,876 to 22,657 (mean 8,887) for the animals that received ITS103.01 alone and from 974 to 11,907 (mean 5,837) for the animals that received ITS103.01 and ITS61.01 together (Fig. 4A and Table S1). During rechallenge, IC50 titers ranged from 3,004 to 39,045 (mean 12,326) for the ITS103.01 animals and from 1,688 to 16,967 (mean 4,374) for the ITS103.01 plus ITS61.01 animals (Fig. 4B and Table S2). IC50 titers correlated well with serum concentrations of ITS103.01 (Fig. S3A-D). ADCC EC50 titers also correlated with serum concentrations of ITS61.01 and ITS103.01 (Fig. S3E, F), consistent with the contribution of both antibodies to this effector function. Notably, neutralization titers in the animals that received ITS103.01, either alone or together with ITS61.01, were 1.5-to 57-fold over the IC50 titer of 685 previously predicted to confer 95% protection against SHIV challenge by passive bnAb transfer (Fig. 4C, D) (*3*). Long-term protection against SIV_mac_239 can therefore be attributed to sustained high level expression of ITS103.01 at concentrations that exceed thresholds necessary to afford 95% protection.

**Fig. 4.**
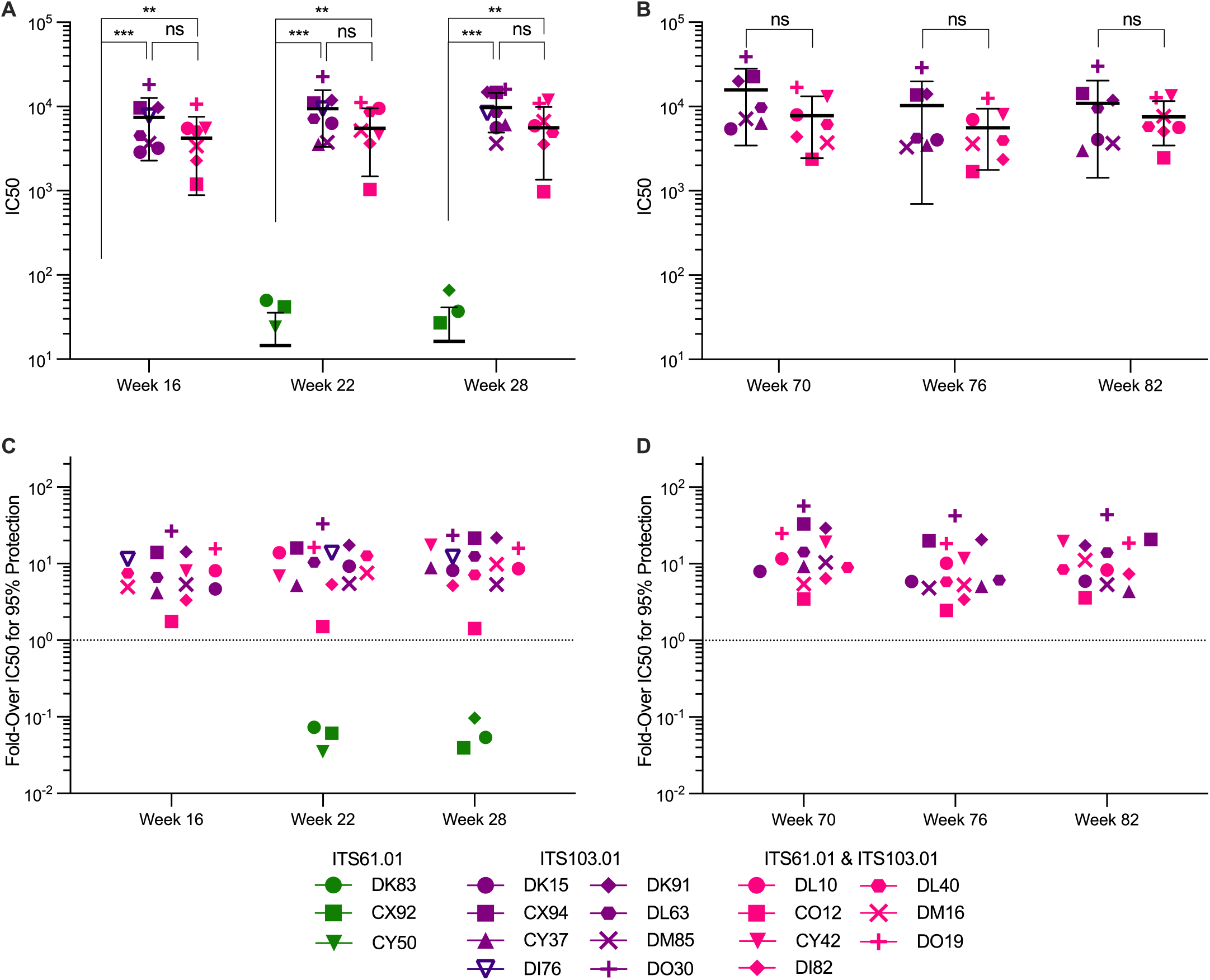
Neutralization titers at the time of SIV_mac_239 challenge. Neutralizing antibody IC50 titers in serum at 16, 22 and 28 weeks (**A**) and at 70, 76 and 82 weeks (**B**) after AAV administration. Error bars indicate standard deviation of the mean for each group. Differences in mean IC50 titers were compared between groups that received vectors encoding ITS61.01, ITS103.01 and ITS61.01 & ITS103.01 using Mann-Whitney tests (‘ns’ not significant, **p<0.005 and ***p<0.0005). (**C, D**) Fold-over the IC50 value estimated to confer 95% protection was calculated by dividing mean IC50 titers for each animal at 16, 22 and 28 weeks (**C**) and at 70, 76 and 82 weeks (**D**) by 685 (*3*). The dotted line corresponds to a value of one.

## DISCUSSION

High and unabated antibody expression was maintained for more than a year in immunocompetent adult rhesus macaques after AAV delivery of antibodies to the SIV Env. Of sixteen animals that received an AAV vector encoding the bnAb ITS103.01, fourteen were completely protected against two rounds of twelve intrarectal SIV_mac_239 challenges more than a year apart. SIV_mac_239 is considered a tier 3, neutralization-resistant virus that is notoriously difficult to protect against by vaccination. Structural studies have revealed especially dense Env glycosylation that make this virus particularly resistant to host antibody responses (*19, 20*).

Indeed, neutralizing antibody titers in animals chronically infected with SIV_mac_239 are typically low and often undetectable (*21, 22*). Recombinant rhesus cytomegalovirus (rhCMV) vectors expressing SIV antigens are the most successful vaccine approach to date, resulting in early containment of SIV_mac_239 in 50-60% of immunized macaques (*23, 24*). Thus, the frequency of complete and durable protection achieved in this study (87.5%) is substantially higher than has been achieved thus far, providing a rigorous demonstration of the potential for AAV delivery of bnAbs to protect against HIV.

Protection can largely if not entirely be attributed to ITS103.01. The IC50 for SIV_mac_239 neutralization by ITS103.01 is remarkably low (∼10 ng/ml) (*14*) and the animals that resisted SIV_mac_239 challenge maintained serum IC50 titers that exceeded the titer predicted to confer 95% protection against SHIV challenge (*3*). Conversely, there was no evidence of protection in the animals that received ITS61.01 alone. This is consistent with a recent study by our group showing that an antibody that mediates potent ADCC in the absence of detectable neutralization is not sufficient for protection (*25*). Although this interpretation is somewhat complicated by high ADA responses in several of the ITS61.01-only animals, no sequence changes in Env were shared among the animals with high levels of ITS61.01 to suggest that this antibody exerted selective pressure on virus replication. While we cannot preclude the possibility that ITS61.01 may have constrained escape from ITS103.01 in animals expressing both antibodies, these results support the idea that neutralization is a critical determinant of antibody-mediated protection.

AAV delivery of naturally isolated rhesus macaque antibodies was key to sustained expression of Env-specific antibodies above protective levels. Previous studies encountered strong ADA responses that prevented vectored antibodies from reaching effective concentrations in most animals. Indeed, 13 of 16 macaques that received vectors encoding “simianized” human HIV bnAbs, in which constant regions of IgG1 were exchanged for their macaque counterparts, and 9 of 12 macaques that received vectors encoding SIV-specific antibodies reconstructed from Fabs isolated by phage display, developed such high ADA responses that serum concentrations of these antibodies plummeted to baseline within a few weeks of AAV administration (*9, 26*). In contrast, ADA responses to ITS61.01 and ITS103.01 were generally low and transient. Only one animal (CY13) developed high ADA titers that curtailed the expression of both antibodies.

Interestingly, ADAs to ITS61.01 were observed in four animals that received the vector encoding this antibody alone but were negligible in the animals that also received the vector for ITS103.01, suggesting that co-expression of ITS103.01 may have inhibited immune responses to ITS61.01.

ITS103.01 is a true bnAb specific for a conserved epitope in the CD4 binding site of SIV gp120 with features characteristic of many HIV bnAbs, including long CDR3H and CDR3L regions generated by extensive somatic hypermutation, which are respectively 22.8% and 8.8% divergent from their predicted germline sequences (*14*). ADA responses to ITS103.01 were nevertheless minimal, resulting in sustained antibody expression in 15 of 16 animals more than a year after AAV delivery with mean serum concentrations in the two treatment groups of 43.7 and 71.6 µg/ml, respectively. Transient immunosuppression with rapamycin prior to and after AAV administration was recently shown to mitigate ADA responses, enabling sustained expression of simianized HIV bnAbs in 12 of 15 macaques (*27*). Although approximately 40% of these animals still developed ADAs and serum concentrations of the delivered antibodies were 2-to 5-fold lower than we observed for ITS103.01 (*27*), these results illustrate the potential for pharmacological suppression of ADAs to enhance antibody expression. Immunomodulatory regimens designed to reduce ADAs may therefore further enhance the frequency of treatment success following AAV delivery of species-matched bnAbs that are naturally less immunogenic.

Infection of one animal despite high serum concentrations of ITS103.01 illustrates the vulnerability of depending on a single bnAb for protection. Sequence analysis of the virus population emerging in this animal (DI76) revealed an Env substitution resulting in the loss of an N-linked glycan previously shown to confer resistance to ITS103.01 (*11*). Although this variant may have been present in the inoculum, low and transient “blips” of viremia have been observed in macaques during serial SIV challenge despite potent bnAbs concentrations many times over levels predicted to prevent infection (*28*). Such subclinical infections, possibly reflecting limited virus replication at local sites of inoculation, create an opportunity for antibody escape. Optimal protection against genetically diverse, naturally transmitted HIV field isolates, will therefore almost certainly require AAV delivery of two or more potent bnAb to prevent the selection of antibody escape variants.

Complete protection in an overwhelming majority of macaques against mucosal SIV_mac_239 challenge more than a year after AAV delivery of a single bnAb illustrates the promise of this approach for achieving long-term protection against HIV. Recombinant AAV has proven safe and effective as a gene therapy platform for the treatment of a variety of hereditary diseases and seven AAV-based therapies have already received approval for use in humans (*29, 30*). Moreover, AAV vectors encoding HIV bnAbs were well tolerated in human volunteers (*31, 32*). Dozens of potent bnAbs targeting diverse epitopes of HIV Env are now available. Thus, durable protection against HIV acquisition may be achieved in people at risk of infection by the careful selection of two or three of these antibodies for AAV delivery.

## MATERIALS AND METHODS

### Study design

The objective of this study was to determine if adeno-associated virus delivery of Env-specific antibodies can protect against mucosal challenge with pathogenic SIV. Rhesus macaques were pre-screened for antibodies to AAV9 and MHC class I genotyped to select thirty-two AAV9-seronegative and *Mamu-B*008-* and *–B*017*-negative animals. Separate groups of eight animals received intramuscular injections of AAV9 vectors encoding the control antibody 17-HD9 (N=8) or the SIV Env-specific antibodies ITS61.01 and ITS103.01, administered alone (N=8 each) or together (N=8). Sixteen weeks later, the animals began a series of twelve weekly low dose intrarectal challenges with SIV_mac_239. Seventy weeks post-AAV administration, the protected animals (N=14) and six naïve control animals (N=6) were rechallenged with second series of twelve weekly low dose inoculations with SIV_mac_239. Peripheral blood was collected at longitudinal time points to monitor viral RNA loads in plasma and antibody concentrations in serum. The number of sampling and experimental replicates are indicated in the figure legends. The number of animals per group was determined from previous experience with SIV_mac_239 infection of rhesus macaques (*25*) and the animals were distributed evenly between experimental and control groups according to sex, age, weight and MHC class I genotype (Table S3).

### Ethics statement

Rhesus macaques (*Macaca mulatta*) were initially housed at BIOQUAL, Inc. (BIOQUAL, Rockville, MD) for AAV administration and for the first round of SIV_mac_239 challenges beginning at week 16 post-AAV. For the second round of SIV_mac_239 challenges beginning at week 70 post-AAV administration, the animals were moved to the Wisconsin National Primate Research Center (WNPRC, Madison, WI). Animals were maintained in accordance with the standards of AAALAC International at BIOQUAL and the WNPRC. Animal experiments were approved by the Institutional Animal Care and Use Committees at BIOQUAL (protocol number 23-119) and at the WNPRC (protocol number G006982) and performed in compliance with the principles described in the Guide for the Care and Use of Laboratory Animals (*33*). Fresh water was always available, commercial monkey chow was provided twice a day, and fresh produce was supplied daily. To minimize any distress, ketamine HCL alone or ketamine HCL in combination with dexmedetomidine were used to sedate animals prior to experimental procedures (e.g., blood collection, AAV administration and SIV infection) and animals were monitored twice a day by animal care and technical staff for signs of morbidity. Analgesics and anti-inflammatories (e.g., buprenorphine, lidocaine and meloxicam) were administered to alleviate pain associated with the experimental procedures. The animals were socially housed in pairs or groups of compatible animals whenever possible.

### Animals

Thirty-eight rhesus macaques of Indian ancestry and free of simian retrovirus type D (SRV), simian T-lymphotropic virus type 1 (STLV-1), SIV and macacine herpesvirus 1 (herpesvirus B) were used for these studies. Animals were pre-screened for AAV9 neutralizing antibodies and were considered negative with no detectable neutralizing activity at a 1:10 dilution of serum (*34*). MHC class I genotyping was performed by sequence-specific PCR in the Sampa Santra Laboratory (Beth Israel Medical Center, Harvard Medical School, Boston, MA). Animals positive for AAV9 neutralizing antibodies or MHC class I alleles associated with spontaneous control of SIV infection (*Mamu-B*008* and *-B*017* (*17, 18*)) were excluded and the experimental and control groups were structured with a similar distribution of male and female animals. The sex, age, weight and MHC class I genotype of each animal is shown in Table S3.

### AAV vector preparation

The AAV transfer plasmids containing AAV2 inverted terminal repeats were previously described (*11*). Codon-optimized (Genscript) sequences for the heavy and light chains of the SIV Env-specific rhesus macaque antibodies ITS61.01 and ITS103.01 (*12, 14*) and the RSV F-specific control antibody 17-HD9 (*13*) were cloned into the AAV transfer plasmid downstream of a CMV enhancer, chicken β-actin promotor and an SV40 intron and upstream of a WPRE and SV40 polyadenylation site. The heavy and light chain reading frames were separated by a P2A ribosomal skip sequence and a furin cleavage site for proteolytic removal of the P2A peptide.

Three tandem repeats of the miRNA binding site miRNA-142T were included in the 3’ UTR to minimize off-target expression in professional antigen presenting cells (*35, 36*). M428L and N434S (LS) substitutions were also introduced into the heavy chain of each antibody to extend their *in vivo* half-lives (*37, 38*).

Recombinant AAV9 vectors were produced as described previously (*39, 40*). HEK293T cells were co-transfected with the AAV transfer plasmid encoding antibody, the pAAV2/9n packaging plasmid expressing Rep2/Cap9 (Addgene, cat. 112865) and the pAdDeltaF6 plasmid expressing adenoviral helper proteins E2a, E4 and VA (Addgene, cat. 112867). AAV9 particles were purified from lysates of transfected cells on CsCl density gradients followed by extensive dialysis against buffer consisting of 5% sorbitol in PBS, after which Pluronic F-68 was added to 0.001% final concentration. Vector concentration in genome copies per ml (gc/ml) was determined by droplet digital PCR for DNAse-resistant templates (*41*). Purity of the final AAV9 preparations was evaluated from a silver-stained SDS-PAGE. For long-term storage, 0.5 ml aliquots of the AAV9 stocks were frozen and stored at −80°C.

### AAV administration

Three separate groups of eight rhesus macaques received AAV9 vectors encoding the SIV Env-specific antibodies ITS61.01 and ITS103.01 individually or together, and eight control animals received an AAV9 vector encoding the RSV F-specific antibody 17-HD9. The AAV9 vectors were administered by intramuscular injection at doses of 3.3×10^12^ gc/kg. To prevent heavy and light chain mixing, vectors encoding ITS61.01 and ITS103.01 were given on opposite sides of the body for the group that received both vectors. To maximize muscle transduction, the AAV9 inoculum was distributed over six separate injection sites on the same side of the body. Each dose of a given vector was suspended in 1 ml of sterile PBS, and 1 ml volumes drawn into tuberculin syringes (Cardinal Health, cat. 309065) were injected into two sites in the quadriceps, two in the deltoids, and two in the biceps.

### SIV challenge

The SIV_mac_239 challenge stock used for this study was prepared at the Wisconsin National Primate Research Center. Supernatant was collected from Vero cells transfected with SIV_mac_239 DNA prepared by ligating *Sph* I-digested plasmids encoding the 5’ and 3’ halves of the viral genome and used to infect CD8-depleted, concanavalin A-activated PBMCs from three cynomolgus macaques. The infected PBMCs were maintained in RPMI medium supplemented with 15% FBS and 50 U/ml IL-2. After re-suspending in fresh medium, the SIV_mac_239 challenge stock (#2282) was harvested the following day (February 12, 2014) by collecting and cryopreserving 0.5 ml aliquots of the cell culture supernatant. The virus concentration of the stock was measured at 192 ng/ml by SIV p27 antigen-capture ELISA and the infectivity titer was determined to be 50,699 TCID_50_/ml by limiting dilution on CEMx174 cells. The identity of the virus stock was confirmed by Illumina MiSeq next generation sequencing as previously described (*25*).

Each of the macaques were challenged weekly by repeated, low dose intrarectal inoculation with SIV_mac_239 for twelve consecutive weeks or until they became infected as determined by detectable viral RNA in plasma. On each day of challenge, a fresh vial of the SIV_mac_239 challenge stock was quickly thawed in a gloved hand, immediately transferred to ice, and diluted to 150 TCID_50_/ml in ice-cold RPMI medium without additives. One ml aliquots of the final virus dilution (150 TCID_50_/ml) were drawn into labeled 3 ml slip-tip syringes for the first round of challenges at BIOQUAL, Inc. and 1 ml syringes for the second round of challenges at the WNPRC. After loading, the syringes were re-capped and stored on ice until inoculation.

Animals were anesthetized (15 mg/kg ketamine-HCl, IM) and positioned with their hindquarters elevated. The needle was removed and the full length of the syringe barrel was slowly inserted into the rectum. The SIV_mac_239 inoculum was gradually expelled, and the syringe was slowly withdrawn, while maintaining the animal’s pelvis in an elevated position. Intrarectal challenges were repeated weekly and blood was collected at each time point to measure viral RNA in plasma. Subsequent challenges were discontinued for infected animals with detectable plasma viral loads.

### Plasma viral load assay

Plasma SIV RNA levels were determined by a two-step real-time qPCR assay. A QIAsymphony SP (Qiagen, Hilden, Germany) automated sample preparation platform along with a Virus/Pathogen DSP midi kit and the *cellfree500* protocol were used to extract viral RNA from 500 µl of rhesus macaque plasma. A reverse primer specific to the *gag* gene of SIV_mac_239 (5’-CAC TAG GTG TCT CTG CAC TAT CTG TTT TG −3’) was annealed to the extracted RNA and then reverse transcribed into cDNA using SuperScript^TM^ III Reverse Transcriptase (Thermo Fisher Scientific, Waltham, MA) along with RNAse Out (Thermo Fisher Scientific, Waltham, MA). The resulting cDNA was treated with RNase H (Thermo Fisher Scientific, Waltham, MA) and then added (2 replicates) to a custom 4x TaqMan™ Gene Expression Master Mix (Thermo Fisher Scientific, Waltham, MA) containing primers and a fluorescently labeled hydrolysis probe specific for the *gag* gene of SIV_mac_239 (forward primer 5’-GTC TGC GTC ATC TGG TGC ATT C −3’, reverse primer 5’-CAC TAG GTG TCT CTG CAC TAT CTG TTT TG −3’, probe 5’- /56-FAM/CTT CCT CAG TGT GTT TCA CTT TCT CTT CTG CG/3BHQ_1/-3’). The qPCR was then carried out on a QuantStudio 3 Real-Time PCR System (Thermo Fisher Scientific, Waltham, MA) using the following thermal cycler parameters: heat to 50°C, hold for 2 min, heat to 95°C, hold for 10 min, then the following parameters are repeated for 50 cycles: heat to 95°C, hold for 15 seconds, cool to 60°C and hold for 1 minute. Mean SIV *gag* RNA copies per reaction were interpolated using quantification cycle data and a serial dilution of a highly characterized custom RNA transcript containing a 730 bp sequence of the SIV *gag* gene. Mean RNA copies per milliliter were then calculated by applying the assay dilution factor (DF=18.72). The limit of quantification (LOQ) for this assay was approximately 62 RNA cp/mL (1.79 log10) with 500 µl of sample.

### ELISA for monitoring monoclonal antibody concentration in serum

ITS61.01 with a C-terminal rhodopsin tag (C9 tag) and untagged ITS103.01 were produced in transfected Expi293 cells (ThermoFisher, cat. A14527) and purified on rProtein A Gravitrap columns (Cytiva Life Sciences, cat. 28985254) according to manufacturer’s suggestions. These antibodies were used to generate a calibration curve in a standard sandwich ELISA at final concentrations ranging from 0.47 to 15 ng/ml.

Reacti-Bind plates (Thermo Scientific, cat. 15041) were coated overnight at 4°C with either 0.8 μg/ml anti-rhodopsin antibody (clone 1D4, MilliporeSigma, cat. MAB5356) or 1.0 μg/ml mouse ITS103.01 anti-idiotype antibody in PBS (*14*). All washes were done with PBS plus 0.07% Tween 20 (PBST) and all incubation steps (blocking, primary and secondary antibody) used PBST containing 5% non-fat dry milk. Experimental sera samples were not heat-inactivated and were applied to the plate as serial two-fold dilutions. Blocking and primary sample incubations were one hour at 37°C and secondary antibody incubations with a 1:6,000 dilution of mouse anti-monkey HRP conjugate (clone SB108a, Southern Biotech, cat. 4700-05) were one hour at 37°C. Bound HRP activity was developed with SureBlue TMB substrate (LGC Clinical Diagnostics, cat. 5120-0075) for 10-15 min, the reaction was stopped with an equal volume of 1 M H_2_S0_4_ and absorbance at 450 nm was quantified on a Victor X4 plate reader (PerkinElmer). Values reported are A_450_ means across all dilutions falling within a valid calibration curve range that are at least two-fold above background and no more than 2.8.

### Anti-drug antibody assay

Anti-drug antibodies were measured by small-scale on-column purification and biotinylation of total IgG from rhesus macaque serum followed by ELISA quantification of the amount of the biotinylated IgG binding to immobilized ITS61.01 or ITS103.01 detected with HRP-conjugated streptavidin exactly as previously described (*11*). For a known negative standard, a sample consisting of equal volumes of pre-immune sera from all experimental animals was used. To anchor the colorimetric responses to a standardized measure, polyclonal rabbit anti-mouse antibody was spiked into pre-immune sera and used as a positive control in the same ELISA but against immobilized total mouse IgG in order to provide a calibration curve. Both types of the controls were treated identically to the experimental samples with respect to purification and biotinylation.

### Neutralization assay

SIV neutralization was measured using a standard TZM-bl assay (*42*) with a few modifications. Serum samples, without heat-inactivation were serially diluted in D10 medium (DMEM containing 10% heat-inactivated FBS) and incubated with SIV_mac_239Δ*vif* in white 96-well plates (Greiner, cat. 82050-736) with 1.5-12 ng of SIV p27 per well, depending on the infectivity of the viral stock used, in a final volume of 100 μl per well. After one hour at 37°C, 15,000 TZM-bl cells (NIH HIV Reagent Program, cat. ARP-8129) of human origin were added in 100 μl of D10 medium. After 72 hours, the medium was carefully aspirated and 100 μl of BriteLite Plus luciferase substrate diluted two-fold in PBS (Revvity, cat. 6066761) was added to each well. The relative light units (RLU) of luciferase activity in each well was measured within 5 min after the addition of substrate using a Victor X4 plate reader (PerkinElmer). After subtracting background luciferase activity in uninfected cells, the luciferase activity was normalized to 100% RLU relative to wells with infected cells without serum or neutralizing antibody. All reported values are means of technical triplicates.

### ADCC assay

ADCC was measured as previously described (*43*) with the following modifications. Five to fifteen million human CEM.NKR-_CCR5_-sLTR-Luc target cells were infected with VSV-G pseudotyped SIVmac239Δ*vif* (85-110 ng p27 per million cells) in 15 ml conical tubes by spinoculation at 1200 *g* in the presence of 40 μg/ml Polybrene for 90 min at 25°C. Three days after infection, the target cells were washed three times and effector cells were washed once with R10 (RPMI 640 medium supplemented with 10% heat-inactivated FBS) and resuspended in assay medium (R10 containing 10 U/ml IL-2). CEM.NKR-_CCR5_-sLTR-Luc target cells (1×10^4^ cells) were mixed with a human NK cell line (KHYG-1 cells) transduced with rhesus macaque CD16 (1×10^5^ cells) in the presence of serial dilutions of serum in a round-bottom 96-well plates (200 μl/well). CEM.NKR-_CCR5_-sLTR-Luc cells and CD16-transduced KHYG-1 cells were established as previously described (*43*) and are available upon request. After an 8-hour incubation at 37°C, 150 μl of the cell suspension from each well was transferred to white 96-well plates (Greiner, cat. 82050-736) containing 100 μl per well of BriteLite Plus luciferase substrate diluted two-fold in PBS. The luciferase activity (RLU) in each well was measured using a Victor X4 plate reader. After correction for the background luciferase activity in wells containing uninfected cells without serum, the measured luciferase activity was normalized to 100% RLU relative to wells with infected cells but without serum. All reported values are means of technical triplicates.

### Antibody concentrations in rectal transudate

Rectal mucosal secretions were collected and processed using Weck-Cel spears (Beaver Visitec, cat. 0008680) with a modification of a previously described protocol (*44*). Each Weck-Cel spear was pre-weighted together with a screw cap 1.5 ml microcentrifuge tube. After wetting the weck with 50 μl of PBS, it was inserted into the rectum and held inside for 5 min. Upon withdrawal, the weck was placed back into the tube and its handle cut off at a pre-determined height so the tube could be capped. The tubes with wecks were kept on ice throughout collection, weighed again and frozen at −80°C for storage before processing. The weight of the cut off handle was determined from averaging 10-20 pieces, a measurement that was found not to exceed a coefficient of variation of 6%.

Upon thawing on ice, the adhered material was manually homogenized with the weck itself in the same tube with 0.3 ml of the freshly prepared ice-cold extraction buffer consisting of 1X PBS, 2 mM EDTA, 0.5% Igepal CA-630, 5 mg/ml BSA, 5 mM sodium azide and 1X protease inhibitors cocktail (ThermoFisher, cat. 87785) and incubated on ice for 15 min. The resulting suspension was transferred to another 1.5 ml microcentrifuge tube with a bottom pinhole made with a 23G needle. The tube was inserted into a 5 ml culture tube (Falcon, cat. 352058) and centrifuged at 2000 g for 5 minutes at 4°C. A second round of homogenization, incubation, and centrifugation was performed with an additional 0.3 ml of extraction buffer, the extract collected in the 5 ml tube was transferred to a 1.5 ml microcentrifuge tube, clarified by centrifugation at 20,000 g for 5 minutes at 4°C. The supernatant was aliquoted, frozen and stored at −80°C for further analysis. Samples with visible blood contamination were excluded from analysis due to the potential contribution of antibodies from plasma.

Monoclonal antibody concentrations in rectal transudates were determined by ELISA using the conditions used to measure antibody concentrations in serum after correction for sample’s dilution after weck pre-wetting and extraction. This weight was assumed to translate to a volume with density of 1 mg/ml. These concentrations were then normalized to total IgG content in the samples to express the amount of recombinant antibody IgG as a percentage of total IgG. In all cases, two rectal weck samples were taken per each time point and the mean values of the replicates are reported.

The ELISA conditions to measure total IgG were as follows. Plates were coated at 4°C overnight with 0.5 μg/ml goat anti-rhesus IgG antibody (SouthernBiotech, cat. 6200-01). Rhesus monkey IgG (SouthernBiotech, cat. 0135-01) was used at concentrations ranging from 3.125 to 100 ng/ml to generate calibration curve. All washes were done with PBST and all incubation steps (blocking, primary and secondary antibody) used PBST containing 5% non-fat dry milk.

Samples were serially diluted two-fold starting at 1:300 and incubated for one hour at 37°C. Bound antibodies were then detected using mouse anti-monkey-HRP (Southern Biotech, cat. 4700-05) at 1:10,000 dilution. Bound HRP activity was visualized with SureBlue TMB substrate (LGC Clinical Diagnostics, cat. 5120-0075) for 10-15 min and the reaction was stopped by adding an equal volume of 1 M H_2_S0_4_ and absorbance at 450 nm was quantified on a Victor X4 plate reader (PerkinElmer). Values reported are A450 means across all dilutions falling within a valid calibration curve range that are at least two-fold above background and no more than 2.8.

### Statistics and data presentation

Statistical methods used in this study include the Mantel-Cox test, Mann-Whitney *U* test and simple linear regression, as indicated in the figure legends. Error bars represent standard deviation of the mean of three or more biological replicates. Significance was determined from unadjusted *p*-values with a detection threshold of 0.05. Nonparametric tests were applied to data that was not normally distributed. All statistical comparisons were performed as two-tailed tests using GraphPad Prism version 8.4.3.

## Author contributions

D.T.E. designed the study. N.M.C., N.K.K., G.N.H., V.A.K., C.M.F., T.N.D. and C.T.D. performed the experiments and analyzed data. H.A., J.T., M.G.L., J.S.H. and S.C. supervised the animal studies. J.X., G.G., M.R.G. and M.R. provided essential reagents. M.R.G., B.F.K., G.G. and D.T.E. supervised laboratory investigations. D.T.E. wrote the manuscript with contributions from all authors.

## Data and materials availability

All data are available in the main text or the supplementary materials. Nucleotide sequences of SIV *env* variants are available at GenBank accession numbers PX921293-PX921619.

## SUPPLEMENTARY MATERIALS

**Fig. S1.**
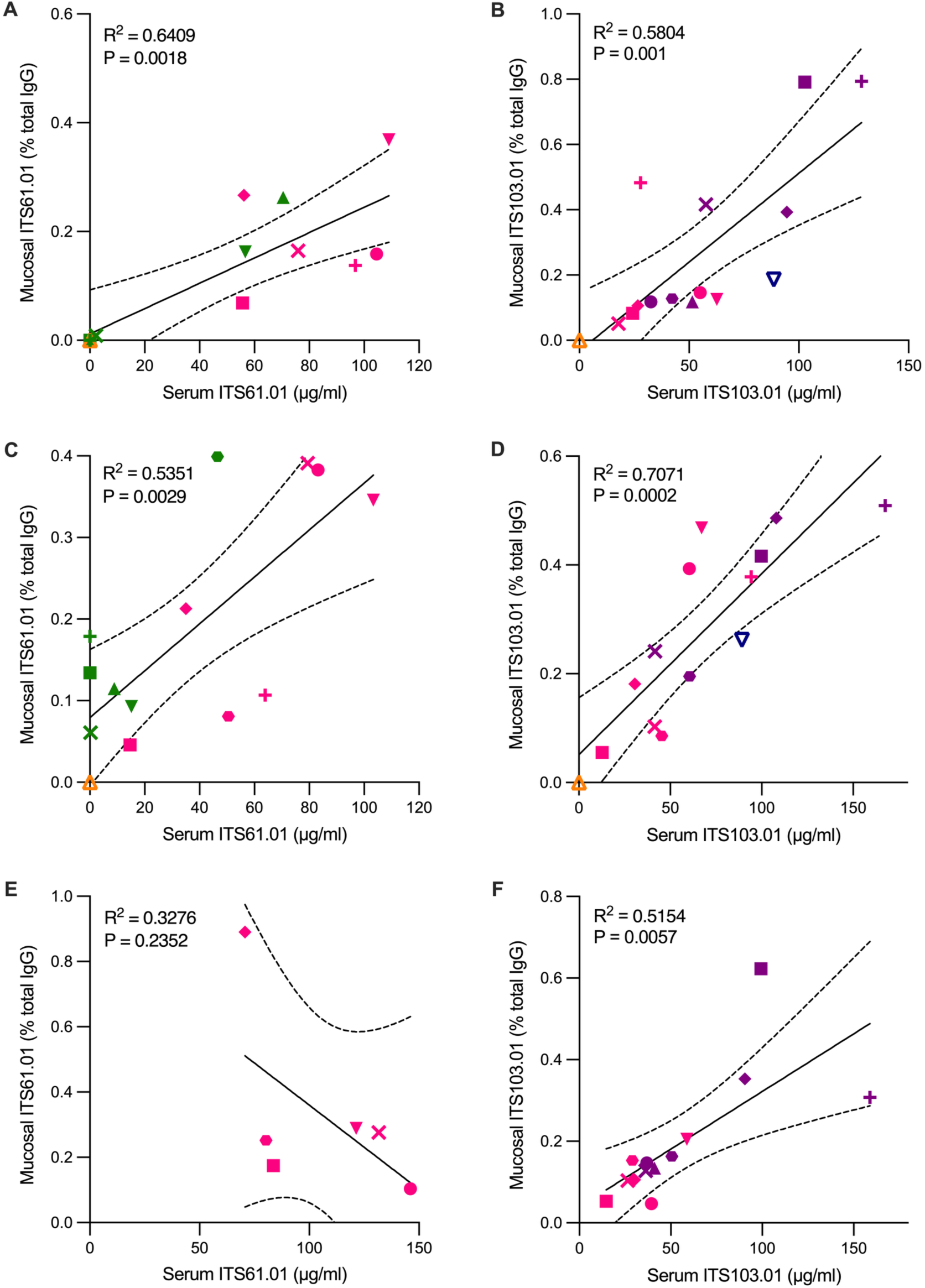
Mucosal antibody concentrations. ITS61.01 and ITS103.01 concentrations as a percentage of total IgG in rectal transudate were compared with serum concentrations of each antibody at week 8 (**A**, **B**), week 12 (**C**, **D**) and week 69 (**E**, **F**) after AAV administration by simple linear regression. The dashed curves indicate 95% confidence intervals.

**Fig. S2.**
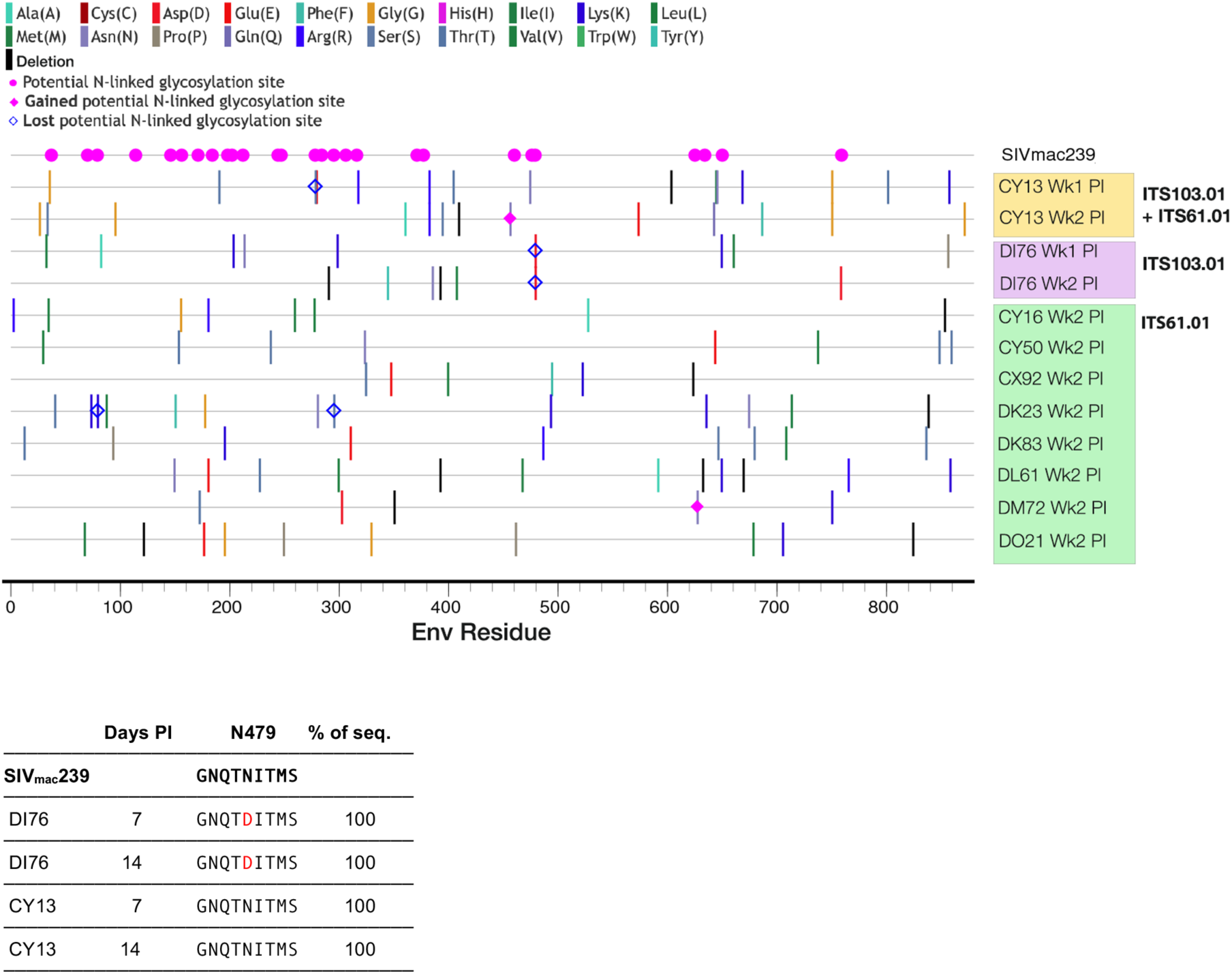
SIV Env substitutions. SIV RNA was isolated from plasma collected at the indicated timepoints after SIV infection. The SIV *env* gene was amplified by RT-PCR after limiting dilution to favor amplification from a single viral genome. At least fifteen independent cDNA products were sequenced at each time point. (**A**) The distribution of amino acid substitutions in Env and the gain or loss of potential N-linked glycosylation sites is summarized by a highlighter plot. (**B**) An asparagine-to-aspartic acid substitution at position 479 (N479D) predicted to result in the loss of N-linked glycosylation of this residue was observed in 100% of sequences on days 7 and 14 post-infection (PI) from animal DI76 but not from CY13.

**Fig. S3.**
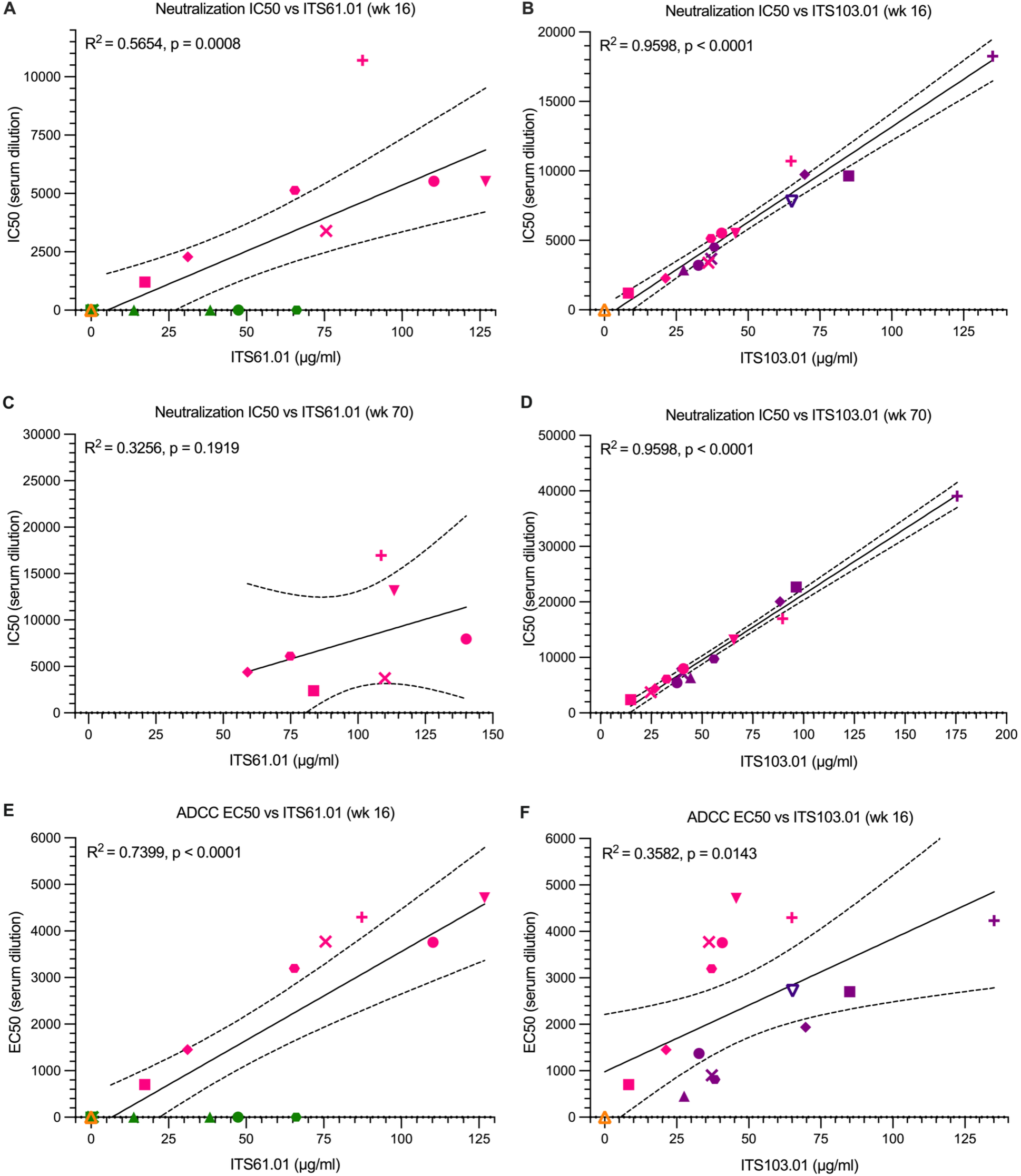
Correlation of neutralization and ADCC with serum concentrations of Env-specific antibodies. Mean neutralization IC50 titers at week 16 (**A**, **B**) and week 70 (**C**, **D**) and mean ADCC EC50 titers at week 16 (**E**, **F**) after AAV administration were compared with mean serum concentrations of ITS61.01 and ITS103.01 at the same timepoints by simple linear regression. The dashed curves indicate 95% confidence intervals.

**Table S1.** Antibody concentrations, neutralization and ADCC during the first series of SIV challenges.

| Animal | ITS61.01 (µg/ml) |  |  | ITS103.01 (µg/ml) |  |  | Neutralization (IC50) |  |  | ADCC (EC50) |
| --- | --- | --- | --- | --- | --- | --- | --- | --- | --- | --- |
|  | Wk 16 | Wk 22 | Wk 28 | Wk 16 | Wk 22 | Wk 28 | Wk 16 | Wk 22 | Wk 28 | Wk 16 |
| DK83 | 47.8±9.3 | 14.3±0.5 | 14.2±1.7 | - | - | - | nd | 50±15 | 37±12 | nd |
| CX92 | nd | nd | nd | - | - | - | nd | 42±20 | 27±14 | nd |
| CY16 | 15.9±4.7 | 9±0.5 | 19.5±1.2 | - | - | - | nd | nd | nd | nd |
| CY50 | 49±15.5 | 16.0±3.0 | 19.7±0.9 | - | - | - | nd | 24±20 | nd | nd |
| DK23 | 0.7±1.2 | nd | nd | - | - | - | nd | nd | 66±35 | nd |
| DL61 | 67.1±15.6 | 41.2±4.2 | 66.8±4.3 | - | - | - | nd | nd | nd | nd |
| DM72 | 0.7±0.5 | 0.2±0.01 | 0.1±0.01 | - | - | - | nd | nd | nd | nd |
| DO21 | 0.4±0.6 | nd | nd | - | - | - | nd | nd | nd | nd |
| DK15 | - | - | - | 32.3±5.3 | 28.8±1.4 | 38.0±1.9 | 3204±892 | 6332±770 | 5607±727 | 1372±422 |
| CX94 | - | - | - | 84.0±19.6 | 80.6±10.6 | 96.1±5.9 | 9635±5632 | 10984±1611 | 14726±1538 | 2700±1243 |
| CY37 | - | - | - | 27.2±5.3 | 27.9±5.3 | 43.1±4 | 2876±275 | 3561±1083 | 6075±1456 | 454±87 |
| DI76 | - | - | - | 64.8±11.3 | 68.4±11.9 | 68.1±12.7 | 7814±1224 | 9417±1872 | 8365±2207 | 2721±1105 |
| DK91 | - | - | - | 68.9±12 | 62.2±2.7 | 101.5±6.7 | 9739±2630 | 11921±1794 | 14864±3432 | 1935±929 |
| DL63 | - | - | - | 37.9±4.9 | 39.1±0.3 | 50.1±5.1 | 4525±1678 | 7156±819 | 8498±3277 | 811±247 |
| DM85 | - | - | - | 37.6±10.6 | 21.5±1.2 | 36.7±1.1 | 3670±1063 | 3757±643 | 3658±224 | 904±42 |
| DO30 | - | - | - | 137.9±16.6 | 131.3±14.5 | 191.3±7.4 | 18252±6053 | 22657±1086 | 16004±2188 | 4232±865 |
| DL10 | 111.8±29.4 | 81.1±2.7 | 107.1±7.5 | 40.3±4.5 | 37.1±1.3 | 42.8±3.9 | 5524±2577 | 9544±4828 | 5903±376 | 3758±3052 |
| CO12 | 20.1±6.5 | 14.8±1.3 | 33.3±1.7 | 8.1±2.1 | 9.0±0.9 | 10.5±0.7 | 1202±389 | 1034±360 | 974±114 | 702±173 |
| CY13 | nd | nd | nd | nd | nd | 1.5±0.3 | nd | nd | nd | nd |
| CY42 | 146.2±50.1 | 93.8±9.4 | 136.0±7.1 | 44.7±10.4 | 55.3±13.3 | 68.7±12.4 | 5516±2384 | 4710±4098 | 11907±645 | 4712±3139 |
| DI82 | 40.9±14.1 | 32±4.3 | 45.9±2.7 | 20.7±3.1 | 15.3±0.3 | 22.4±2.1 | 2281±474 | 3657±1634 | 3559±1151 | 1452±724 |
| DL40 | 67.4±10 | 46±3.8 | 59.5±3.2 | 36.8±7.7 | 31.7±0.7 | 36.2±0.7 | 5138±2175 | 8695±3419 | 4882±1525 | 3195±2349 |
| DM16 | 75.2±16.8 | 71.7±7.1 | 94.6±5.9 | 36.6±6.7 | 28.4±3.9 | 38.7±4.8 | 3399±1619 | 5201±1366 | 6704±1025 | 3771±2027 |
| DO19 | 80.4±24.6 | 63.7±0.8 | 103.0±5 | 65.8±9.7 | 62.2±2.6 | 87.9±7.7 | 10703±1190 | 11146±1763 | 10901±3116 | 4298±2014 |

Rhesus macaques and their corresponding data are shaded according to treatment group; ITS61.01 (green), ITS103.01 (purple) and ITS61.01 & ITS103.01 (rose). Neutralization IC50 and ADCC EC50 titers are expressed as fold dilution of serum. Means and standard deviations were calculated from three independent assays and 'nd' indicates not detectable.

**Table S2.** Antibody concentrations, neutralization and ADCC during SIV rechallenge.

| Animal | ITS61.01 (µg/ml) |  |  | ITS103.01 (µg/ml) |  |  | Neutralization (IC50) |  |  |
| --- | --- | --- | --- | --- | --- | --- | --- | --- | --- |
|  | Wk 70 | Wk 76 | Wk 82 | Wk 70 | Wk 76 | Wk 82 | Wk 70 | Wk 76 | Wk 82 |
| DK15 | - | - | - | 37.7±5.1 | 32±2 | 35.6±4.6 | 5443±2584 | 4040±519 | 4060±1700 |
| CX94 | - | - | - | 96.4±15.2 | 82.9±2.9 | 89.1±8.3 | 22705±5138 | 13705±2849 | 14285±3179 |
| CY37 | - | - | - | 44.3±6.4 | 37±3.9 | 26.9±2.5 | 6360±1931 | 3476±938 | 3004±439 |
| DK91 | - | - | - | 88.4±8.6 | 89±8.2 | 80.2±6.6 | 20066±3222 | 14097±3744 | 11848±3558 |
| DL63 | - | - | - | 55.9±9.3 | 46.1±4.8 | 53.6±6.1 | 9733±1407 | 4202±386 | 9571±2838 |
| DM85 | - | - | - | 40.4±3.9 | 32.5±2.3 | 33.4±2.7 | 7164±505 | 3308±289 | 3676±1153 |
| DO30 | - | - | - | 175.7±10 | 182.6±18.5 | 167.2±12 | 39045±10054 | 29041±8309 | 30009±4047 |
| DL10 | 140.1±10.1 | 105.4±6.5 | 122.1±9.7 | 40.8±4.2 | 42±2.3 | 37.6±4.8 | 7967±2828 | 6988±747 | 5663±1272 |
| CO12 | 83.6±2.4 | 90.6±3.7 | 81.6±2 | 14.9±1.8 | 15±0.9 | 16.3±1.3 | 2380±747 | 1688±69 | 2472±760 |
| CY42 | 113.4±1.2 | 90.6±5.1 | 113.3±5.5 | 65.7±9.9 | 63.3±6.6 | 67.1±6.9 | 13152±2869 | 8019±1303 | 13486±6266 |
| DI82 | 59±2 | 46.8±1.2 | 55±3.8 | 36.9±3.3 | 27.7±3 | 26.1±2.8 | 4396±1387 | 2348±38 | 5073±2786 |
| DL40 | 74.8±3.6 | 65.6±4.7 | 85.9±3.6 | 32.5±4.4 | 28.2±3 | 33.8±1.1 | 6121±870 | 3977±201 | 5760±2750 |
| DM16 | 110±6.8 | 90.6±2.3 | 110.3±6.1 | 24.7±2.6 | 38.5±4.3 | 43±3.6 | 3730±826 | 3630±292 | 7659±4561 |
| DO19 | 108.6±5.4 | 87.6±9.7 | 110.9±6.3 | 89.7±8.6 | 92.6±8.1 | 85.4±8.4 | 16967±6581 | 12520±2035 | 12744±2068 |

Rhesus macaques and their corresponding data are shaded according to treatment group; ITS103.01 (purple) and ITS61.01 & ITS103.01 (rose). Neutralization IC50 and ADCC EC50 titers are expressed as fold dilution of serum. Means and standard deviations were calculated from three independent assays.

**Table S3.** Rhesus macaques.

| Animal | Sex | Age (yrs) | Weight (kg) | MHC Class I Genotype |
| --- | --- | --- | --- | --- |
| CH56 | female | 9.63 | 6.36 | Mamu-A1*001-pos, -B*008-neg, -B*017-neg |
| DK17 | male | 3.12 | 4.58 | Mamu-A1*001-neg, -B*008-neg, -B*017-neg |
| DK44 | male | 2.96 | 3.78 | Mamu-A1*001-neg, -B*008-neg, -B*017-neg |
| DK59 | male | 3.07 | 4.02 | Mamu-A1*001-neg, -B*008-neg, -B*017-neg |
| DL06 | male | 2.59 | 3.62 | Mamu-A1*001-neg, -B*008-neg, -B*017-neg |
| DL58 | male | 3.00 | 3.88 | Mamu-A1*001-neg, -B*008-neg, -B*017-neg |
| CI17 | female | 9.15 | 6.04 | Mamu-A1*001-pos, -B*008-neg, -B*017-neg |
| CP29 | female | 7.29 | 7.54 | Mamu-A1*001-pos, -B*008-neg, -B*017-neg |
| DK83 | male | 3.07 | 4.04 | Mamu-A1*001-neg, -B*008-neg, -B*017-neg |
| CX92 | male | 3.13 | 4.68 | Mamu-A1*001-neg, -B*008-neg, -B*017-neg |
| CY16 | male | 3.15 | 4.32 | Mamu-A1*001-neg, -B*008-neg, -B*017-neg |
| CY50 | male | 3.06 | 4.46 | Mamu-A1*001-neg, -B*008-neg, -B*017-neg |
| DK23 | male | 2.99 | 3.84 | Mamu-A1*001-neg, -B*008-neg, -B*017-neg |
| DL61 | male | 3.06 | 4.02 | Mamu-A1*001-neg, -B*008-neg, -B*017-neg |
| DM72 | male | 3.04 | 4.28 | Mamu-A1*001-pos, -B*008-neg, -B*017-neg |
| DO21 | male | 2.99 | 4.40 | Mamu-A1*001-neg, -B*008-neg, -B*017-neg |
| DK15 | male | 3.05 | 4.36 | Mamu-A1*001-neg, -B*008-neg, -B*017-neg |
| CX94 | male | 3.21 | 4.86 | Mamu-A1*001-neg, -B*008-neg, -B*017-neg |
| CY37 | male | 2.95 | 4.40 | Mamu-A1*001-neg, -B*008-neg, -B*017-neg |
| DI76 | male | 3.28 | 3.94 | Mamu-A1*001-neg, -B*008-neg, -B*017-neg |
| DK91 | male | 3.31 | 4.88 | Mamu-A1*001-neg, -B*008-neg, -B*017-neg |
| DL63 | male | 3.07 | 4.44 | Mamu-A1*001-neg, -B*008-neg, -B*017-neg |
| DM85 | male | 3.07 | 4.04 | Mamu-A1*001-neg, -B*008-neg, -B*017-neg |
| DO30 | male | 2.67 | 3.70 | Mamu-A1*001-neg, -B*008-neg, -B*017-neg |
| DL10 | male | 2.69 | 3.84 | Mamu-A1*001-neg, -B*008-neg, -B*017-neg |
| CO12 | female | 7.87 | 5.98 | Mamu-A1*001-pos, -B*008-neg, -B*017-neg |
| CY13 | male | 3.19 | 4.14 | Mamu-A1*001-neg, -B*008-neg, -B*017-neg |
| CY42 | male | 3.00 | 4.66 | Mamu-A1*001-neg, -B*008-neg, -B*017-neg |
| DI82 | male | 3.19 | 3.76 | Mamu-A1*001-neg, -B*008-neg, -B*017-neg |
| DL40 | male | 2.75 | 4.14 | Mamu-A1*001-neg, -B*008-neg, -B*017-neg |
| DM16 | male | 3.03 | 5.30 | Mamu-A1*001-neg, -B*008-neg, -B*017-neg |
| DO19 | male | 2.79 | 4.24 | Mamu-A1*001-neg, -B*008-neg, -B*017-neg |
| r10054 | female | 15.03 | 7.91 | Mamu-A1*001-pos, -B*008-neg, -B*017-neg |
| r20056 | male | 5.14 | 7.48 | Mamu-A1*001-neg, -B*008-neg, -B*017-neg |
| r21011 | male | 3.41 | 6.72 | Mamu-A1*001-neg, -B*008-neg, -B*017-neg |
| r22054 | male | 2.09 | 5.63 | Mamu-A1*001-neg, -B*008-neg, -B*017-neg |
| r22095 | male | 2.64 | 4.39 | Mamu-A1*001-neg, -B*008-neg, -B*017-neg |
| r23021 | male | 2.33 | 4.33 | Mamu-A1*001-neg, -B*008-neg, -B*017-neg |

Rhesus macaques and corresponding data are shaded according to treatment group; 17-HD9 (blue), ITS61.01 (green), ITS103.01 (purple), ITS61.01 & ITS103.01 (rose) and naïve controls (grey). Ages and weights are at the time of the first SIV challenge.

